# Human photoreceptors switch from autonomous axon extension to cell-mediated process pulling during synaptic marker redistribution

**DOI:** 10.1101/2021.10.10.463810

**Authors:** Sarah K. Rempel, Madalynn J. Welch, Allison L. Ludwig, M. Joseph Phillips, Yochana Kancherla, Donald J. Zack, David M. Gamm, Timothy M. Gomez

**Affiliations:** Department of Neuroscience, University of Wisconsin – Madison, WI 53706; Department of Ophthalmology and Visual Sciences, University of Wisconsin – Madison, Madison, WI, 53705; McPherson Eye Research Institute, University of Wisconsin – Madison, Madison, WI 53706; Waisman Center, University of Wisconsin – Madison, Madison, WI 53705; Department of Ophthalmology, Johns Hopkins University, Baltimore, MD 21287

**Keywords:** retinal development, axon extension, regeneration, photoreceptor migration

## Abstract

Photoreceptors (PRs) are the primary visual sensory cells, and their loss leads to blindness that is currently incurable. Cell replacement therapy holds promise as a therapeutic approach to restore vision to those who have lost PRs through damage or disease. While PR transplant research is ongoing in animal models, success is hindered by our limited understanding of PR axon growth during development and regeneration. Using a human pluripotent stem cell (hPSC) reporter line that labels PRs (WA09 CRX^+/tdTomato^), we generated retinal organoids in order to study mechanisms of PR process extension. We found that the earliest born PRs exhibit autonomous axon extension from dynamic terminals that appear similar to projection neuron growth cones. However, as hPSC-derived PRs age from 40 to 80 days of differentiation, they lose dynamic terminals in 2D plated cultures and within 3D retinal organoids, which does not correlate with cell birth date. Using a rod-specific hPSC reporter line (WA09 NRL^+/eGFP^), we further determined that rod PRs never form motile growth cones. Interestingly, PRs without motile terminals are still capable of extending axons, but neurites are generated from process stretching via their attachment to motile non-PR cells, which underlies the observed differences in PR neurite lengths on different substrata. While immobile PR terminals express actin, it is less polymerized and less organized than actin present in motile terminals. However, immobile PRs do localize synaptic proteins to their terminals, suggesting a normal developmental progression. These findings help inform the development of PR transplant therapies to treat blinding diseases and provide a platform to test treatments that restore autonomous PR axon extension.

**Significance Statement:** Loss of photoreceptors (PRs) in the retina through damage or disease causes irreversible vision loss and blindness. One treatment approach is to replace lost cells with transplanted human stem cell-derived PRs, but this requires PR axons to integrate into the host retina to restore the required neural connections. For this strategy to succeed, we need to understand how PRs extend processes to their targets during development *in situ*, and whether dissociated human stem cell (hPSC)-derived PRs behave in a similar fashion. In this paper, we show that hPSC-PRs have only a short window during which they are capable of autonomous axon extension, which has implications for PR transplant efforts and for our basic understanding of human retinal development.

## Introduction

Retinal photoreceptors (PRs) are primary visual sensory neurons, and therefore their proper connection and function is critical for vision. Loss of PRs due to disease or damage results in vision loss and irreversible blindness, and one potential treatment approach is cell replacement therapy with human pluripotent stem cell (hPSC)-derived PRs (1–4). While PR transplant research is ongoing (5–14), our limited understanding of key aspects of normal PR development – including axonal process initiation, targeting, axon extension and guidance, and synaptic maturation – is hindering successful transplantation.

While some aspects of retinal development are well understood, relatively little is known about the mechanisms controlling extension of mammalian PR axons. Ramón y Cajal first described the basics of early PR morphological development more than one hundred years ago, detailing three major stages in sections of pre- and post-natal mammalian retinas (15). As live cells were not directly observed, the dynamics of axon extension and retraction could only be speculated by Cajal. His observations were largely corroborated by electron microscopy of developing mouse retina sections (16). Stage one, the germinal stage, is characterized by the presence of mitotic, roughly spherical, apically-located retinal progenitor cells (RPCs). In stage 2, RPCs begin to exit the cell cycle, and newly born PRs transition to the unipolar stage, in which a thin apical process connects to the outer surface of the neural retina, with a small protuberance that will be the future outer segment. Lastly, PRs enter the bipolar stage, where they extend a process tipped with a growth cone into the intermediate layers that eventually retracts to reside in the newly formed outer plexiform layer (OPL). Research using a ferret model showed that developing rod PR processes initially target one of two distinct layers of the inner plexiform layer (IPL), past PRs’ normal target layer (the OPL), where they express synaptic markers and co-stratify with cholinergic amacrine cells (17, 18). Cones over-projecting to the IPL in fetal human and non-human primate retina have been observed in immunostained sections (19, 20). While much has been gleaned from fixed tissues, this research is limited in its observations of static images rather than dynamic, living cells.

A major technical challenge to achieving functional PR replacement is the need to promote connections between donor PR axons and host inner retina neuron (i.e. bipolar cell) dendrites. To this end, it would be helpful to understand the intrinsic and extrinsic factors that inhibit or promote PR axon growth. While research on PR axon development is limited, a pair of studies by Kljavin *et al*. examined dissociated rodent retinal cells in fixed cultures and found that rod axons were longest when co-cultured with the dominant glial cell type in the retina, Müller glia (MG), suggesting that rods extend axons along MG (8, 21).

While little is directly known about PR axon growth, we know quite a lot about the general mechanisms driving axon extension by projection neurons, such as retinal ganglion cells (RGCs) (22). Canonical axon outgrowth occurs through autonomous activity of a dynamic terminal growth cone, which mechanically pulls itself across a semi-rigid adhesive substratum through membrane and cytoskeletal remodeling, followed by axon consolidation at its trailing edge (23). In this way, growth cones are progressively and autonomously guided through their environment while the axon is assembled and maintained distally behind it, leading to persistent elongation. Different types of neurons are known to respond to distinct environmental cues through established regulatory pathways that either promote or inhibit the elongation of axons by modifying their membranes and cytoskeleton (22, 24, 25).

Based on these prior studies of RGCs, we initially hypothesized that PR axon extension would occur via mechanisms similar to those of long projection neurons. However, in 2D cultures of dissociated hPSC-derived retinal organoids (ROs), we discovered that while such mechanisms are required for long-projecting axons to navigate across diverse tissues and make complex guidance decisions, they are not strictly utilized for short-projecting, specialized neurons like PRs. Using live imaging to observe hPSC-PR maturation in real time, we found that they have a short developmental window in which traditional dynamic growth cones that facilitate canonical axon extension are present. Thereafter, these terminals become static, and by day 80 of differentiation, very few PRs are capable of autonomous axon growth. Only early PRs, excluding rods, have dynamic growth cones and autonomous axon growth capability. Interestingly, within that early population, the presence of a dynamic terminal does not appear to be dependent upon cell birth date. In addition, many of the PR axons observed in heterogenous, dissociated RO cultures result from non-autonomous stretching by motile non-PR cells. The age-related reduction in dynamic PR terminals correlates with a reduction in F-actin, and also with an increase in localization of Synaptic Vesicle Protein 2 and Synaptotagmin-1 to the PR terminals. Importantly, the decrease in dynamic PR terminals observed in 2D cultures is also seen over the same time period using live imaging of intact 3D ROs.

## Results

### Photoreceptors lose autonomous terminal motility within weeks after being generated

We performed time-lapse live cell imaging of hPSC-PRs dissociated from ROs on day 80 of differentiation (d80) to test whether their axons extend in a similar manner to projection neurons. This timepoint was initially chosen for analysis because it is within the age range often used for transplantation experiments (14, 26). We used ROs differentiated from a WA09 CRX^+/tdTomato^ reporter line, which robustly labels all post-mitotic PRs (27). ROs derived from human PSCs have been extensively studied and are known to form all neuroretinal cell types, closely recapitulating normal human developmental timelines for marker expression and cellular morphology and developing organized retinal features such as lamination (28–31). In these ROs, the earliest-born PRs (presumptive cones) begin to appear around day 30, while rod PRs are first born at approximately day 70 (27).

Surprisingly, live cell imaging of PRs from ROs dissociated at d80 revealed that PR axon terminals, which appeared to be morphologically similar to traditional growth cones, were largely immobile and had little to no autonomous movement (Fig. 1A, C). A more thorough characterization of PR terminals across early differentiation ages revealed that at very early ages, some post-mitotic PRs had dynamic terminals capable of displacement like classic projection neuron growth cones (Fig. 1A). To quantify terminal dynamics, we measured the change in thresholded area of terminals normalized to terminal size, where *Dynamicity* = ([*Area of Protrusion*] + [*Area of Retraction*])/[*Total Area of Terminal*], averaged across each frame of the time series. We observed a decrease in dynamicity and terminal centroid displacement from d40 to d80 on PDL-Laminin (PDL-LN) substrata (Fig. 1B-C).

**Figure 1.**
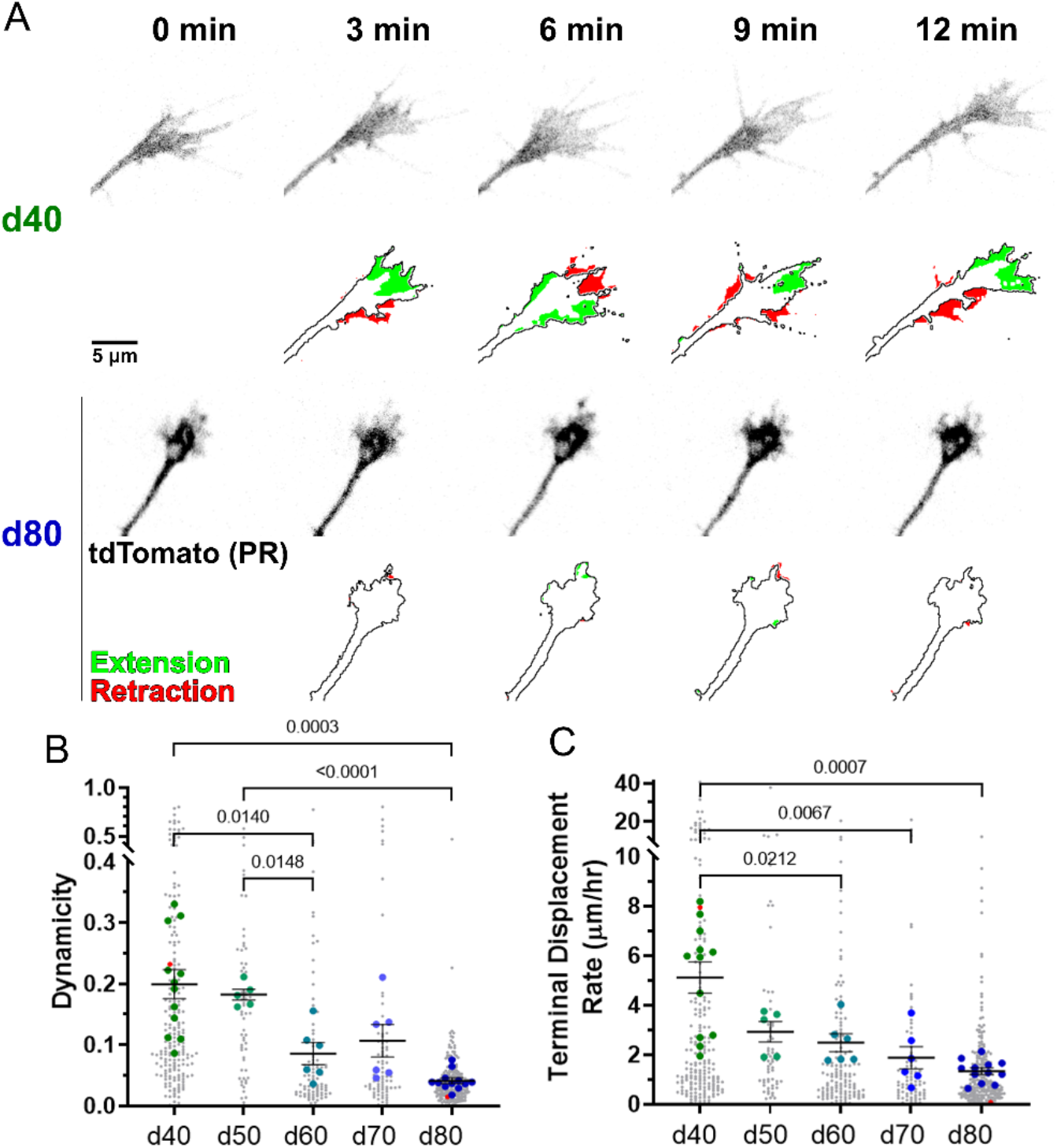
Dynamic day 40 PRs growth cones capable of autonomous displacement become immobile by day 80. **A:** Top rows show tdTomato-labeled PR terminals, while bottom rows show the PR terminal outlined in black, with the areas extended and retracted between time points shown in green and red respectively, for PRs dissociated on d40 (top section) and d80 (bottom section). **B:** Dynamicity ([Area of Protrusion + Area of Retraction]/[Terminal Area]) decreases with age on both PDL and PDL-LN. Red points correspond to the cells shown in A. **C:** Displacement Rate of PR terminals decreases with age on both PDL and PDL-LN. Red points correspond to the cells shown in A. For B and C, statistics were performed on the mean value per dish, indicated by large colored dots, and individual PR values are presented in grey to show population range and variance. Mean and SEM per dish are shown. N=5-12 dishes/condition with 6-38 PRs/dish. P-values <0.05 shown for comparisons from Brown-Forsythe and Welch ANOVA with Dunnett’s T3 multiple comparison test.

To determine whether age-dependent terminal dynamicity was intrinsic or due to signals from aging non-PR cells in the heterogenous RO cultures, we co-cultured old and young PRs together. d40 WA09-CRX^+/tdtomato^ RO cells were cultured with unlabelled d80 WA09 RO cells, or d80 PR-labeled RO cells with d40 unlabeled RO cells, to determine how dynamicity of PRs are affected by RO cell age. We found that d40 PRs cultured with d80 RO cells were much more dynamic than d80 PRs cultured with d40 RO cells, and the dynamicity of PRs was similar to the same ages from homochronic cultures, suggesting that the age-related decrease in PR terminal dynamicity is intrinsic (Supplemental Fig. 1). Interference reflection microscopy (IRM), which indicates how closely associated terminals are to the substratum, revealed that the stationary terminals of d80 PRs are more adherent than are the motile terminals of d40 PRs (Supplemental Fig. 2) (32, 33). Together, these results suggest that an intrinsic change occurs within PR cells with age causing increased terminal stability.

**Figure 2.**
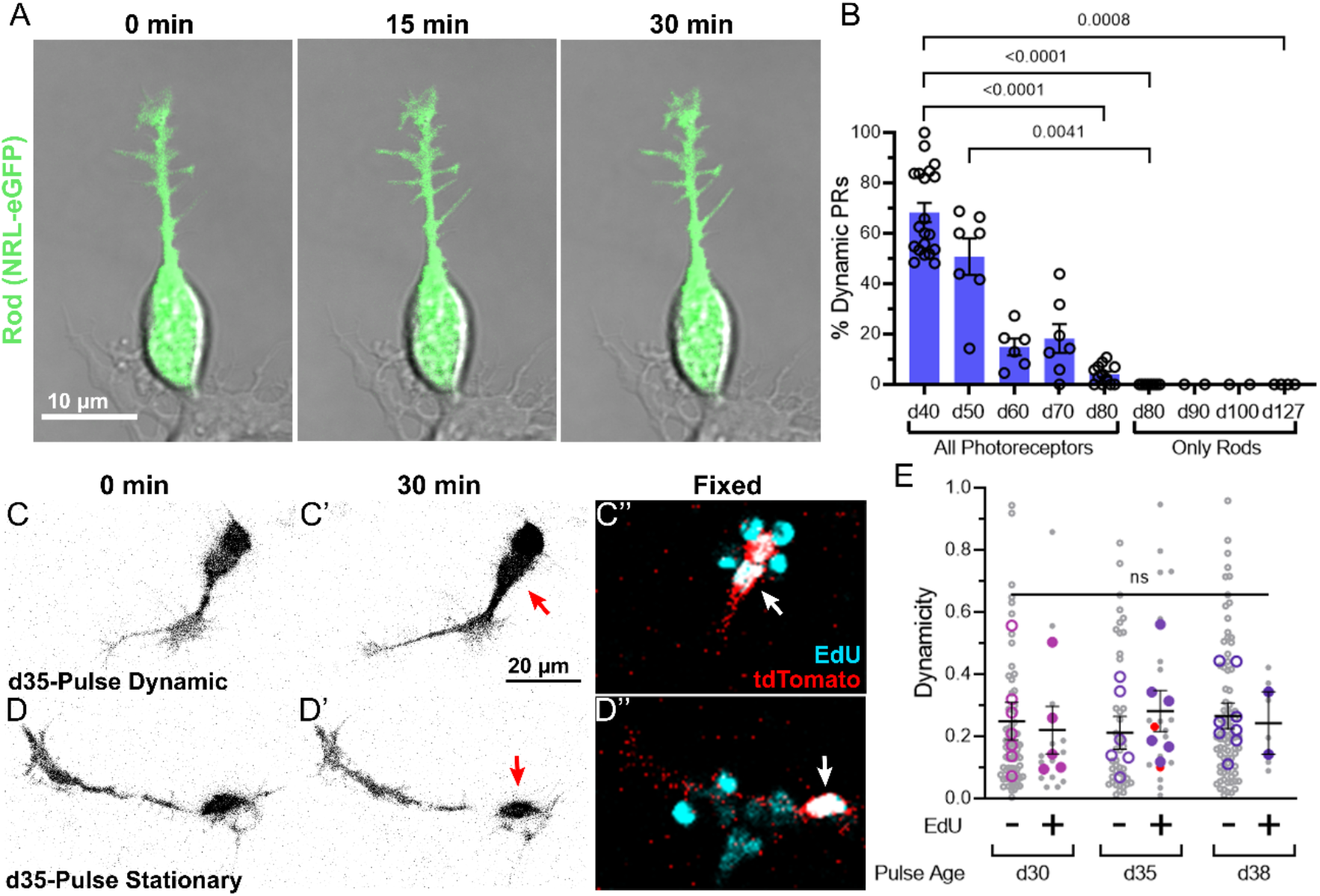
Age-dependent loss of terminal motility occurs in presumptive cone PRs independent of birth date, while rod terminals never appear dynamic. **A:** Time-lapse imaging of a d80 dissociated rod PR (green) shows a static terminal over 30min. **B:** Qualitative assessment of dynamicity shows a decrease in proportion of PRs with dynamic terminals for all PRs (from dissociations of WA09 CRX^+/tdTomato^ ROs) over d40-80, while rods only (from dissociations of WA09 NRL^+/eGFP^ ROs) never exhibit dynamic terminals between d80-127. N=2-20 dishes/age with 5-36 PRs/dish and 21-120 PRs/age. Each data point is the percent of PRs with a dynamic terminal per dish, averaged between assessments from two independent analysts. Mean and SEM shown. P-values <0.05 shown for comparisons from Kruskal-Wallis with Dunn’s multiple comparison test. **C**,**D:** Some d40 PRs that are EdU+ (C’, at most 5 days post terminal division upon dissociation) continue to exhibit dynamic terminal movements during live time-lapse imaging (C), while other d40 PRs that are EdU+ (D’) have stationary terminals (D). Arrows indicate cell bodies of tdTomato+/EdU+ cells. **E:** Dynamicity ([Area of Protrusion + Area of Retraction]/[Terminal Area]) of d40 PRs does not correlate with birth date, as PRs labeled with EdU at different ages had similar dynamicity. N=2-8 dishes/pulse age and EdU category with 2-18 PRs/dish. Red points correspond to the cells shown in C and D. No significant differences in means per two-way ANOVA with Šídák’s multiple comparison test (p>0.99 for all). Mean and SEM are shown.

### Photoreceptor terminal dynamicity is subtype dependent

The age-related decline in dynamic PR terminals coincided with the onset of rod PR birth, suggesting two independent, but potentially related, hypotheses. First, early born, cone PR terminals may be more dynamic than rod terminals, and the age-related decrease in observed dynamicity is simply due to the PR population shifting from being all new cones to a mixture of cones and rods. Second, recently born PRs may be more likely to have dynamic terminals, and over time, newborn PRs make up a shrinking proportion of the total PR population. To test the first hypothesis, we used a rod-specific reporter line (NRL^+/eGFP^) created on the same WA09 background as the CRX^+/tdTomato^ pan-PR reporter line (34). ROs derived from the WA09 NRL^+/eGFP^ line were dissociated at d80, shortly after onset of NRL expression, and at d90, d100, and d127. The time span (47 days) of investigation corresponds roughly to the time span examined using the WA09 CRX^+/tdTomato^ pan-PR reporter line (40 days).

To assess rod PR dynamicity, we used a qualitative classification analysis, since the lower eGFP signal in rod terminals was insufficient to conduct the same quantitative analysis shown in Figure 1. Two independent and masked-to-condition analysts scored each PR as “dynamic” or “stationary”, and the percent of dynamic PRs in each dish was calculated and averaged between analysts. Dynamic terminals were defined as those with at least one period of extension that was not directly influenced by physical interaction with another cell. To validate the qualitative assessment, we first compared it to the earlier quantitative analyses performed using time-lapse sequences of d40-80 WA09 CRX^+/tdTomato^ PRs. As in the automated analysis, the proportion of dynamic CRX^+/tdTomato^ PRs decreased with age from d40 to d80 (Fig. 2B). Following validation of the analytic methods, we examined PR time-lapse sequences from d80-127 rods, and none possessed dynamic terminals (n >600 rods) (Fig. 2B).

To test the second hypothesis — that recently born PRs have dynamic terminals that become more stationary with time after birth — we pulsed WA09 CRX^+/tdTomato^ ROs with thymidine analog 5-ethynyl-2’-deoxyuridine (EdU) at d30, d35, or d38 to identify newly divided cells. Treated ROs were dissociated at d40, imaged live, and terminal dynamicity was quantitatively analyzed as described above. Cultures were fixed after live imaging and stained for EdU to correlate birth date with PR terminal dynamicity (Fig. 2C). If recent birth date were positively correlated with increased terminal motility, we would expect that PRs labeled with EdU in the more distant past would have more stationary terminals compared to PRs labeled by a recent pulse. On the contrary, we observed no relationship between birth date and PR dynamicity; all cultures had EdU+ and EdU-PRs with similar ranges of terminal dynamicity (Fig. 2C-E).

### Photoreceptors undergo axon elongation through non-cell autonomous pulling

Despite the absence of dynamic growth cones capable of autonomous movement in d80 PRs, average PR neurite length was similar between d40 and d80 PRs (Fig. 3A,B). If d80 PRs do not undergo growth cone-mediated extension, what mechanism drives axon elongation? Closer observation revealed that PRs had longer neurites in 2D cultures when they were associated with other cells (Fig. 3A,B). Examination of long-term time-lapse imaging led us to suspect that non-cell autonomous pulling by motile non-PR cells drives hPSC-PR extension at this stage of development (Supplemental Fig. 3).

**Figure 3.**
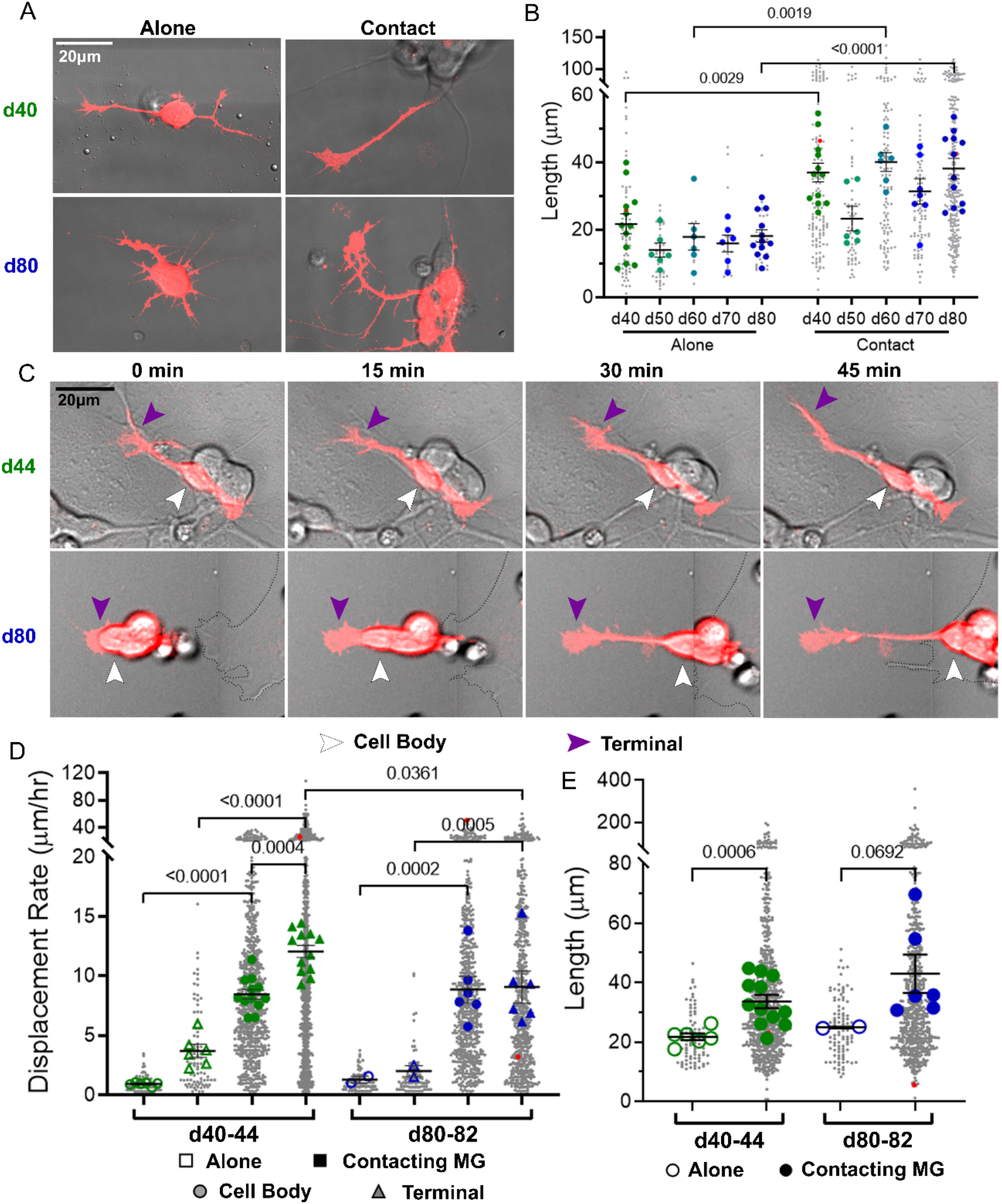
Day 80 PR axon elongation occurs through non-cell autonomous process stretching mediated by co-cultured motile cells. **A**,**B:** PRs contacting other cells in RO culture have longer neurites compared to isolated PRs at the start of imaging. Note axon length also does not change with age. N=3-12 dishes/condition with 6-36 PRs/dish. P-values <0.05 shown for comparisons from two-way ANOVA with Šídák’s multiple comparison test. Red points in B correspond to the cells shown in A. **C:** In MG co-culture with RO cells, a motile d44 PR terminal (purple arrowheads) contributes to axon elongation with little cell body movement (white arrowhead), while a static d80 PR terminal emerges as the cell body is pulled rearward by MG cell (dotted line). **D:** Terminal displacement of d40-44 PRs is greater than cell body displacement on and off MG, while d80-82 PRs have equal terminal and cell body displacement rates while contacting MG, with little movement in isolation. Red points correspond to the cells shown in C. P-values <0.05 shown for comparisons from three-way ANOVA with Šídák’s multiple comparison test. **E:** Total axon lengths of PRs of any age at the start of imaging are shorter in isolated PRs compared to MG-contacting PRs. Red points correspond to the cells shown in C. P-values <0.05 shown for comparisons from Brown-Forsythe and Welch ANOVA with Dunnett’s T3 multiple comparison test. For B, D, and E, large colored shapes show mean value per dish while smaller gray dots show value for individual PRs. Mean and SEM per dish shown. N=2-6 dishes/condition with 6-136 PRs/dish.

Previous studies have shown that Müller glia (MG) co-culture enhanced neurite extension of rods isolated from post-natal rodents retinas (8, 21), and these effects were hypothesized to be due to growth-promoting signals from MG which in turn enhanced PR axon extension. To explore whether this phenomenon also occurs in hPSC-PR cultures, we co-cultured dissociated d40-44 and d80-82 WA09 CRX^+/tdTomato^ ROs with ImM10 cells, an immortalized murine MG line (35) (Fig. 3C). We then measured terminal and cell body displacement of PRs from time-lapse movies, categorizing them as either contacting or not contacting MG during the duration of imaging. Terminal displacement greater than cell body displacement would be consistent with growth cone-mediated cell autonomous extension, while equal terminal and cell body displacements would be consistent with non-cell autonomous pulling by one or more associated cells. We found that both terminal and cell body displacement increased substantially with MG contact in both d40-44 and d80-82 PRs, suggesting MG pulling that was indiscriminate with respect to cell compartment at both ages (Fig. 3D). However, displacements were consistent with some cell autonomous axon extension for d40 PRs, but suggested only non-cell autonomous pulling for d80 PRs (Fig. 3D). Indeed, we observed cases of *de novo* axon generation when PRs that were initially without neurites had their cell bodies carried by motile MG, resulting in the generation of an extended neurite connecting the displaced PR cell body with its stationary, adherent terminal (Fig. 3C). For PRs not contacting MG, cell body displacement was low for both d40 and d80 PRs, while terminal displacement was higher than cell body displacement for d40 PRs, but not d80 PRs (Fig. 3D). Additionally, this type of extension appears to be passive stretching rather than active extension based on the thinning of the axon that takes place when a PR cell body is rapidly moved away from an immobile substratum-bound terminal (Supplemental Movie 1). We conclude that MG enhance PR axon extension at studied ages by pulling PR terminals or cell bodies, rather than by promoting cell autonomous growth cone-mediated PR axon extension through alternative means such as factor secretion.

While it appears that d80 hPSC-PRs depend on motile non-PRs for non-cell autonomous neurite stretching (Fig. 3), it is possible that autonomous terminal motility is substratum-specific *in vivo*. Cell adhesion molecules (CAMs) and extracellular matrix (ECM) proteins regulate axon extension of multiple neuronal types through direct effects on growth cone motility (33, 36–40). Previous research also suggests that PR axon extension similarly depends on the substratum, with MG cells providing the most robust support in fixed co-culture analysis (21, 38). Therefore, we tested the effects of several different substrata on autonomous and cell-contact-mediated PR process extension: Laminin, Neural Cell Adhesion Molecule (NCAM), and Neural-cadherin (Ncad). We found that culture substrata do influence PR axon length, but only for PRs in contact with other cells, suggesting the culture substrata do not affect PR process extension directly, but only indirectly through effects on the non-PR cell population (Fig. 4). Three-way ANOVA of dish averages showed large contributions to variation from contact with other cells (35.1% of variation, p<0.0001) and substrata (16.9%, p<0.0001), with a much smaller contribution of age (0.94%, p=0.003). Interactions between substrata and contact (8.7%, p<0.0001) and substrata and age (3.7%, p<0.0001) also contributed to observed variance, while variance attributable to interactions between age and contact were negligible (0.05%, p=0.507).

**Figure 4.**
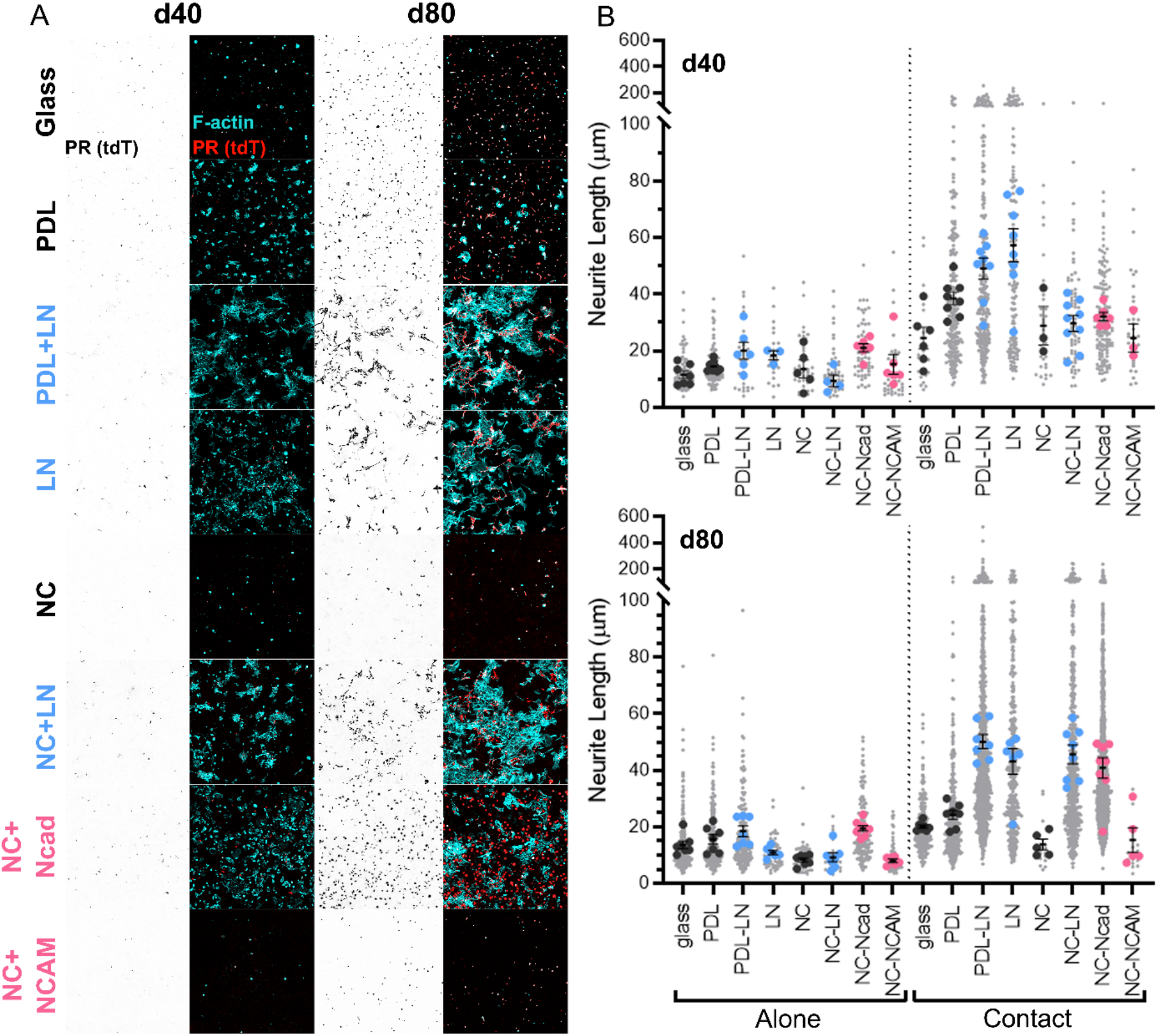
Axon length is not substratum-dependent for isolated d40 or d80 PRs. **A:** d40 or d80 ROs were dissociated and cultured on uncoated glass, poly-*D*-lysine (PDL), PDL plus Laminin (PDL-LN), LN alone, nitrocellulose (NC), NC-LN, NC-Neural-cadherin (NC-Ncad), or NC-Neural Cell Adhesion Molecule (NC-NCAM), and stained for F-actin with phalloidin (cyan) to visualize all cells in the cultures. **B:** PR neurite lengths were similar on all substrata for d40 and d80 PRs in isolation (left). Differences in neurite lengths between substrata are only observed in d40 and d80 PRs that were contacting other cells at the time of fixation (right). Larger colored dots indicate mean value per well, and small gray dots indicate values for individual PRs. Mean and SEM per dish are shown. N=3-9 dishes/condition with 2-454 PRs/dish.

### Photoreceptor terminals lose F-actin with developmental age

The loss of motile terminals with developmental age in the PR population suggests a change in cytoskeletal dynamics, as the cytoskeleton drives cell morphology and motility. The dominant cytoskeletal component controlling canonical growth cone morphology is filamentous-actin (F-actin) (22, 23). Monomeric G-actin polymerizes to form F-actin, which classically organizes into a branched network within lamellipodia and parallel bundles within filopodia. This classical organization is observed across species and neuron types (39, 41, 42).

Using super-resolution Stimulated Emission Depletion (STED) fluorescence microscopy of dye-conjugated phalloidin to label F-actin, we compared F-actin organization in the terminals of d40 PRs, d80 PRs, and d46 retinal ganglion cells RGCs, which were also differentiated from WA09 CRX^+/tdTomato^ and were identified by axon length. RGC growth cones exhibited classic F-actin organization, with intensely labelled bundles of F-actin extending into filopodia, and finer, ordered F-actin mesh throughout the lamellipodia (Fig. 5A). Some d40 PR terminals showed classic F-actin organization (Fig. 5A’), while others had no bright filopodial F-actin bundles and no ordered mesh (Fig. 5A’’). F-actin in d80 PR terminals always appeared disorganized, with dim filaments arranged throughout the terminal and often a few light puncta (Fig. 5A’’’).

**Figure 5.**
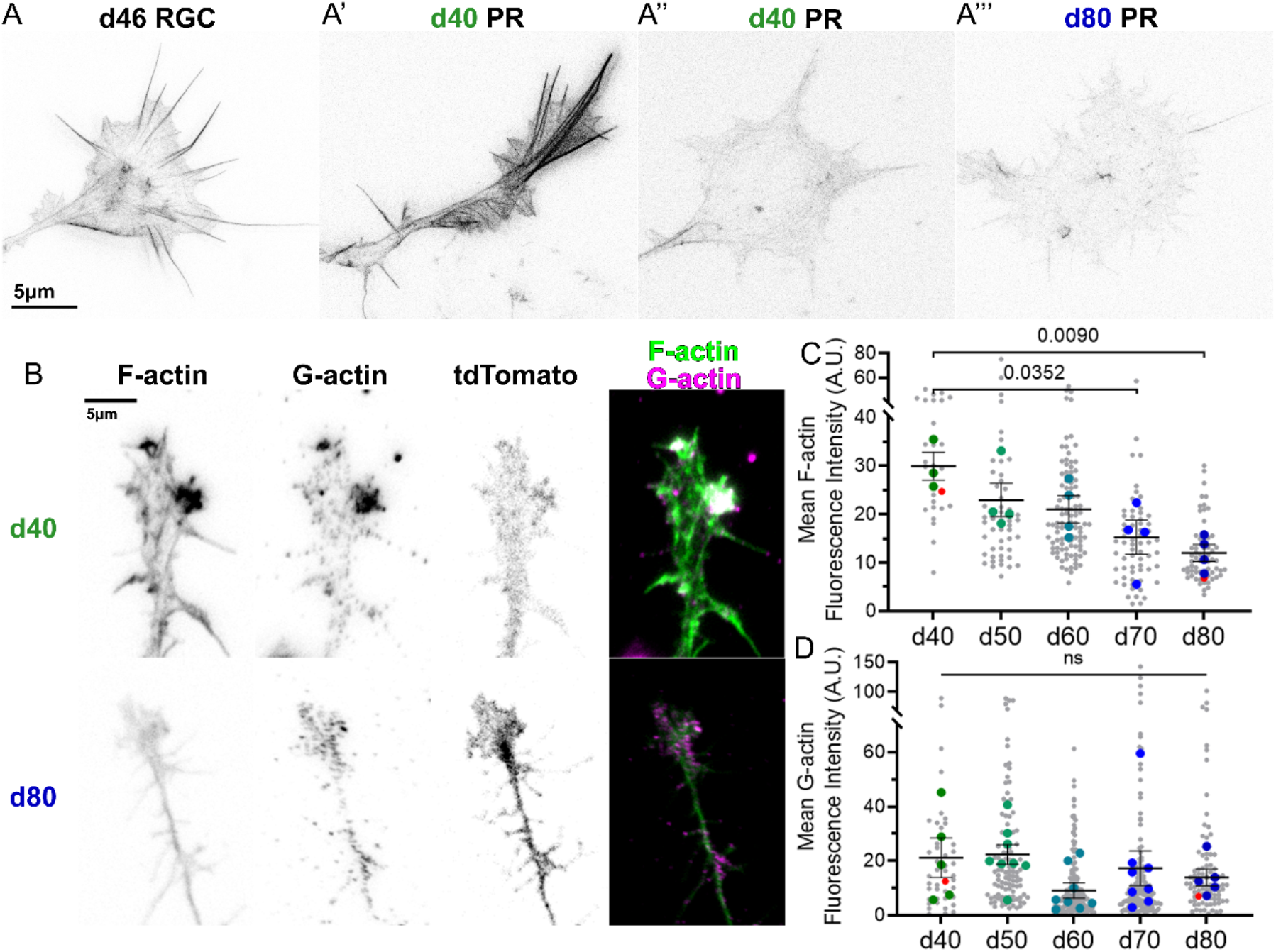
F-actin content and organization decreases in PR terminals with age. **A:** Super-resolution STED inverse contrast images of a d46 RGC growth cone, two d40 PR terminals, and a d80 PR terminal. **B:** Inverse contrast images of d40 and d80 PRs stained with phalloidin to visualize F-actin (green in merge) and immunostained for G-actin (magenta in merge). **C:** F-actin fluorescence intensity decreases with PR age from d40-d80. Red points correspond to the cells shown in B. P-values <0.05 shown for comparisons from one-way ANOVA with Tukey’s multiple comparison test. **D:** G-actin fluorescence intensity is not significantly different as PRs age. Red points correspond to the cells shown in B. P-values <0.05 shown for comparisons from Kruskal-Wallis with Dunn’s multiple comparison test. For C and D, large colored dots show average value per dish while smaller gray dots show values of individual PRs. Mean and SEM per dish are shown. Statistics were performed on dish averages. N=3-8 dishes/age with 6-29 PRs/dish.

Using confocal microscopy, we quantified the fluorescence intensity of F-actin staining together with G-actin immunolabeling (43). The overall F-actin intensity within PR terminals decreased with PR age from d40 to d80, but G-actin intensity did not change significantly with age (Fig. 5C,D). Because F-actin is formed from pools of monomeric G-actin, this finding suggests that while the building blocks of growth cones are expressed in PRs, F-actin is not polymerizing or organizing in some d40 PRs and in nearly all d80 PRs, possibly due to differences in the signaling pathways regulating F-actin assembly.

### Photoreceptors polarize with age and localize synaptic markers to their terminals

Typically, synaptic markers become localized to terminals following cessation of axon extension. To assess whether this phenomenon also occurs in dissociated hPSC-PRs, we immunostained d40-45 and d80-84 PRs for Synaptic Vesicle Protein 2 A (SV2) and Synaptotagmin-1 (Syt-1), two presynaptic markers that are expressed early in PR development (19, 44, 45). We found a striking change in SV2 localization between these ages, with SV2 in d40-45 PRs often localizing to the cell body, near the axon initial segment (Fig. 6A). However, by day 80-84, SV2 labeling shifted to neurites and axon terminals (Fig. 6A). Accordingly, the ratio of SV2 fluorescence intensity in the neurite relative to the cell body was higher in the d80-84 PRs compared to d40-45 PRs. This change in ratio was not due to differences in volume of the neurites and cell bodies, as the mean fluorescence intensity ratio of these two compartments of the tdTomato signal was similar between ages (d40-45 mean neurite/cell body ratio=0.38, S.D.=0.04; d80-84 mean neurite/cell body ratio=0.49, S.D.=0.06). Syt-1 staining also showed changes in localization between d40 and d80 PRs with stronger localization of Syt-1 to the neurites of d80 PR terminals (Fig. 6C,D). Again, the tdTomato ratio between neurites and cell bodies suggested that differences in volume were not the source of differences in Syt-1 fluorescence intensity ratios (d40 mean=0.59, S.D.=0.07; d80 mean=0.51, S.D.=0.03). These results suggest that the immobile terminals of older PRs are being converted to a distinct compartment of the cell, with polarized trafficking of synaptic components to their adherent terminals.

**Figure 6.**
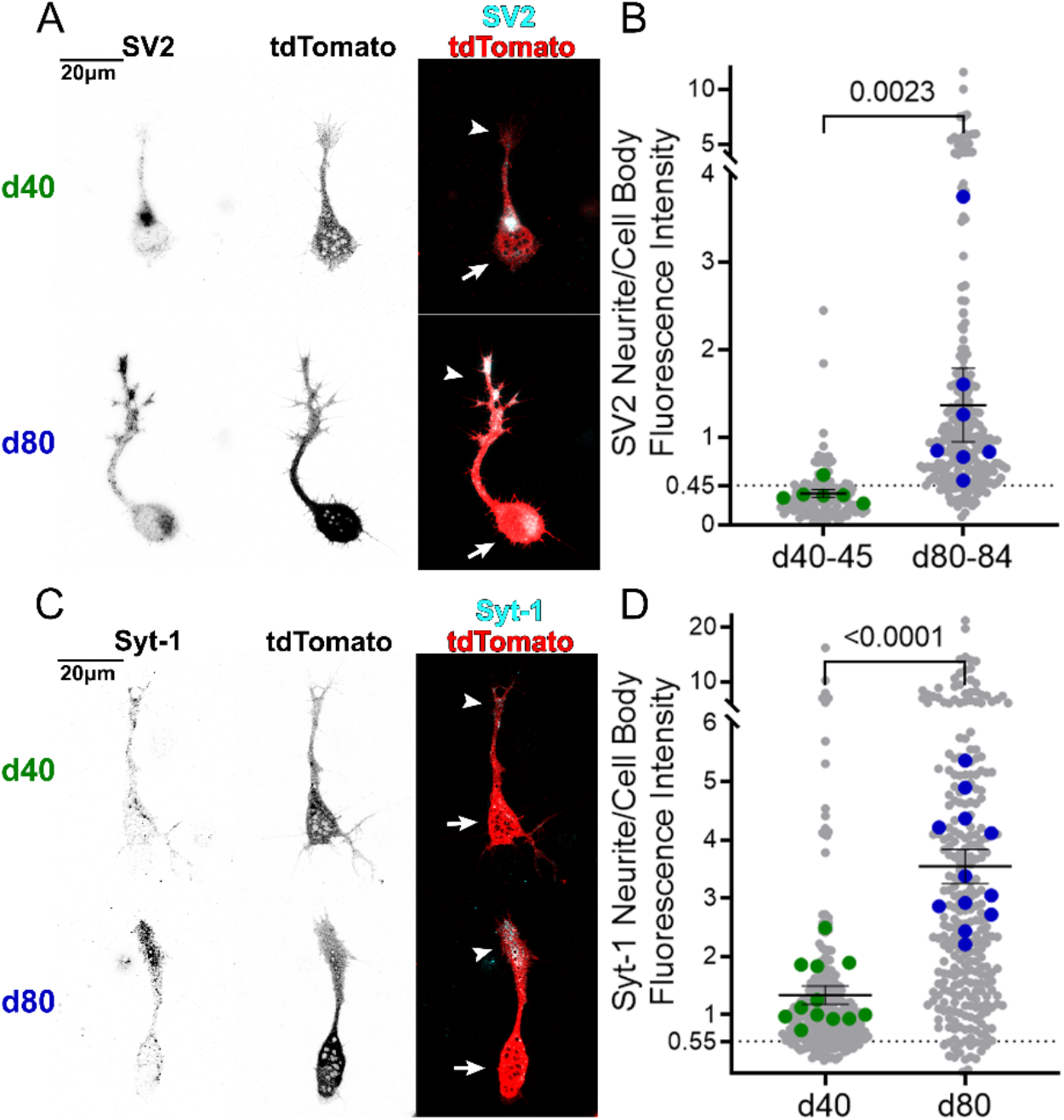
PRs localize synaptic proteins to their terminals as they age. **A:** Synaptic marker Synaptic Vesicle Protein 2 (SV2) localizes to the cell body of a d40 PR, but targets to the terminal of a d80 PRs. **B:** The fluorescence intensity ratio of SV2 immunostaining in neurites divided by cell bodies is higher in d80-84 PRs than d40-45 PRs. Red points correspond to the cells shown in A. N=6-7 dishes/age with 11-76 PRs/dish. P-value shown for Mann-Whitney test. **C:** Synaptic marker Synaptotagmin-1 (Syt1) is localized more strongly to d80 PR terminals than it is to d40 PR terminals. **D:** The fluorescence intensity ratio of Syt1 immunostaining in neurites divided by cell bodies is higher in d80 PRs than d40 PRs. Red points correspond to the cells shown in C. P-value shown for Mann-Whitney test. For B and D, dotted line indicates mean value of tdTomato Neurite/Cell Body Fluorescence Intensity ratio to show the baseline for volume fill. Larger colored dots show mean values per dish while smaller gray dots show values for individual PRs. Statistics were performed on dish averages. N=12 dishes/age with 10-29 PRs/dish. Mean and SEM per dish are shown. Arrows indicate cell bodies and arrowheads indicate terminals.

### Photoreceptors within retinal organoids migrate, have dynamic growth cone-like terminals, and become less dynamic with age

While the use of dissociated 2D cultures simplifies examination of PR axon extension, it does not recapitulate the 3D environment in which PRs develop or into which they will ultimately be transplanted. To assess PR morphology and behavior across ages within the three-dimensional RO microenvironment, we used two-photon microscopy to image PRs live within intact ROs at d40, d60, and d80. PRs, like all retinal cells in these ROs and consistent with normal developing retina, are born on the apical surface (28). However, most PRs at d40 are located basally within the hPSC-ROs. By d60, many PRs have migrated back to the apical surface, but some remain basally located, while by d80 almost all PRs are found near the apical surface (Fig. 7A). To quantify this migration, we measured the ratio of the mean tdTomato fluorescence intensity of the outer half of each RO to the inner half and found that this ratio increases dramatically with age. Additionally, the percentage of PRs within each RO that have an apical neurite decreases with age, concurrent with an increase in the percentage of PRs that have a basal neurite. Finally, at d40 we observe many PRs with what appear to be autonomously dynamic terminals, based on their movement relative to surrounding landmarks and rapid extension and retraction of filopodial and lamellipodial protrusions lateral to the direction of growth (Fig. 7E). In contrast, by d80 most PRs instead have stationary terminals (Fig. 7E,F) and little change in the sum of the length of their neurites compared to dynamic PRs at d40 and d60 (Fig. 7G).

**Figure 7.**
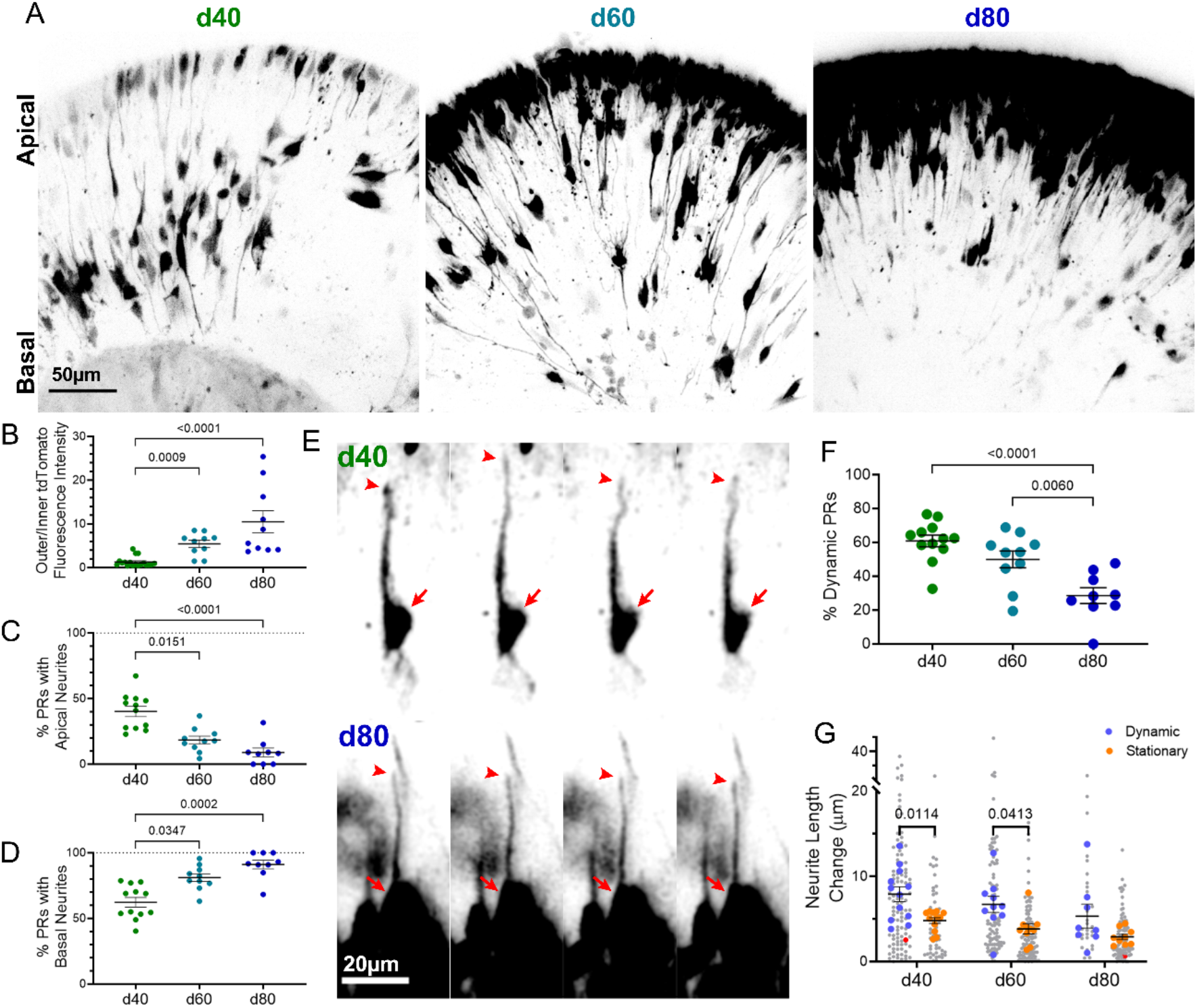
PRs migrate and have dynamic growth cones within ROs that become less motile with age. **A:** Maximum projections of tdTomato signal within 50 μm optical sections of d40, d60, or d80 ROs. **B:** Outer half to inner half ratio of mean tdTomato fluorescence intensity of RO increases with age, reflecting migration of tdTomato+ PRs from the interior of the RO to the apical surface from d40 to d80. N=10-18 ROs/age. P-values <0.05 shown for comparisons from Kruskal-Wallis with Dunn’s multiple comparisons test. **C**,**D:** The percentage of PRs within each RO with an apical neurite decreases with age (C), while the percentage with a basal neurite increases with age (D), reflecting the transition to a unipolar population of PRs with a basal neurite. N= 9-12 ROs/age with 5-31 PRs/RO. P-values <0.05 shown for comparisons from Kruskal-Wallis with Dunn’s multiple comparisons test. **E:** Time-lapse images of a d40 PR with dynamic terminal movement and a d80 PR with a stationary terminal. Arrows indicate cell bodies and arrowheads indicate terminals. **F:** The percentage of PRs within each RO that have a dynamic terminal decreases with age. N= 9-12 ROs/age with 5-31 PRs/RO. P-values shown for comparisons from one-way ANOVA with Tukey’s multiple comparisons test. **G:** PRs with dynamic terminals have greater total neurite length changes (e.g. change in the sum of all neurite lengths) compared to PRs with stationary terminals, and length changes of dynamic terminals decrease with age. Red points correspond to the cells shown in E. N=8-12 ROs/age with 5-31 PRs/RO. P-values <0.05 shown for comparisons from two-way ANOVA with Šídák’s multiple comparison test. For B-D, F, G, mean and SEM per RO are shown.

## Discussion

Here we report that hPSC-PRs generated within ROs have a short window during which they can autonomously extend axons, starting within 10 days after the beginning of the cone birth period and ending 6 weeks later. On the other hand, rod PRs never exhibit dynamic growth cones that support autonomous axon extension at the times we studied. However, PRs incapable of autonomous growth can still generate neurites in culture, and this is accomplished through pulling by motile non-PR cells, such as MG. The loss of PR motility we observe is associated with decreased F-actin and increased synaptic marker expression within stable, maturing PR terminals. Finally, we observe similar behavior by PRs within 3D ROs.

The loss of PR terminal motility correlates with a reduction in F-actin, but not G actin (Fig. 5), suggesting that signals promoting cytoskeletal polymerization are lost within PRs as they age. While prior reports have examined the role of cytoskeletal regulators in injured or degenerating adult PRs (46–49), we are not aware of any previous research on cytoskeletal regulators in developing mammalian PR axons. Together with our results showing delivery of synaptic components to older PR terminals (Fig. 6), these results indicate that PRs are shifting from an early outgrowth phase to a synaptogenesis phase as they mature. Further research is necessary to understand what molecular changes are responsible for motility changes in PR terminals, which may lead to strategies to enhance motility of donor hPSC-PRs post-transplantation.

Interestingly, we found that even PRs incapable of cell autonomous axon extension were able to extend axons when pulled by a motile non-PR cell (Fig. 3). Previous work has shown that axons can be stretched to great lengths by experimental manipulations (50) and axons clearly lengthen due to body growth (51, 52). However, to our knowledge this is the first report showing that direct cell-cell interactions are responsible for axon lengthening. This important finding may help inform PR transplant research, as motile cells and processes within the degenerating retina, such as reactive MG, microglia, or possibly even dendrites from bipolar cells or horizontal cells, could physically stretch neurites of transplanted PRs. Additionally, it is notable that previous reports of conditions that promote PR neurite extension have universally found that the effects are dependent on the presence of other cells (8, 21, 53), consistent with the non-autonomous pulling mechanism shown here.

Fixed human and non-human primate fetal retinal histology shows that PR axon projection to the IPL occurs early in development (19, 20). This understudied phenomenon was explored most fully in ferret, where it was found that PRs extend long processes beyond their target layer to the IPL to make bi-stratified putative synaptic contacts before retracting those processes to ultimately reside in the OPL to begin synaptogenesis (17, 18). We found evidence of PR migration within ROs and observed examples of dynamic growth cone-like structures on young PR terminals that could relate to this early developmental overprojection. Similar basal migration of hPSC-PRs in ROs has been observed previously using an alternative differentiation protocol (54), suggesting this is a general feature of human ROs rather than a culture method-specific phenomenon. Intriguingly, recent research on rod migration in mouse retina has found that PRs undergo passive basal migration primarily caused by apical proliferation, and they must undergo active dynein-mediated apical migration to remain within the ONL (55). Together with our findings, this suggests the intriguing hypothesis that rod axon formation could result from adhesion of the rod terminal within the future OPL followed by apical migration of the cell body away from the terminal.

These findings have important implications for retinal therapies directed at treating blindness by transplanting hPSC-derived PRs into damaged or degenerated retina. Our results suggest that hPSC-PRs may not be universally competent to autonomously extend axons towards dendritic partners. Instead, strategies may need to focus on augmenting bipolar cell dendrite growth toward transplanted donor PRs.

While human fetal samples, *in vivo* animal models, and *in vitro* models have all played a role in furthering our understanding of human retinal development, the experiments presented here demonstrate that hPSC-derived cultures can provide unique insight into the dynamics of human retinogenesis. Our study also underscores the capabilities and limitations of hPSC-PRs, which can help inform cell replacement strategies designed to restore vision in patients with PR loss.

## Materials and Methods

### Retinal Organoid Differentiation

Pluripotent stem cells (WA09 CRX^+/tdTomato^ and WA09 NRL^+/eGFP^) were maintained on Matrigel (Corning #354230) in mTeSR1 (STEMCELL Technologies #85850) or StemFlex (Gibco A3349401) and passaged with Versene upon reaching 60-90% confluency. Differentiation to retinal organoids was based on published protocols (56) with minor modifications and began by lifting embryoid bodies with 2mg/mL Dispase II protease (Sigma #D4693) in DMEM/F12 (Gibco #11330-032) and suspending in stem cell media. Over the next 4 days, embryoid bodies were transitioned to Neural Induction Medium (NIM) [DMEM/F12 (Gibco #11330-032), 1x N2 Supplement (Gibco #17502-048), 1x MEM non-essential amino acids (Gibco #11140-050), 2ug/mL Heparin (STEMCELL Technologies #07980), 1x (2mM) Glutamax (Gibco #35050-061)]. At day 7, embryoid bodies were plated on Matrigel (0.17mg/mL in DMEM/F12) in NIM with 10uM BMP-4 (R&D Systems #314BP010). At day 16, media was changed to Retinal Differentiation Media (RDM) [7:3 DMEM (Gibco 11965-118):F12 (Gibco #11765-054), 1x B-27 supplement (Gibco #17504-044), 1x Antibiotic-Antimycotic (Gibco #15240-062), and 2-10% FBS (Peak Serum, Inc. #PS-FBS)]. From day 7 until dissection, half media changes were performed every 1-3 days. At day 24-30, organoids were manually dissected. The following day, retinal organoids (identified by their phase bright, laminar appearance under a light microscope) were manually separated and transferred to a flask coated with poly-HEMA (Sigma #P3932-10G) to prevent adhesion. Flasks of suspended retinal organoids were fed fresh media every 3-4 days. See Supplemental Table 1 for number of biological replicates (differentiations) and technical replicates (dishes, wells, or ROs) in each experiment. ROs that were used for dissociation experiments were differentiated in RDM with 2% FBS, and ROs that were used for whole RO live imaging were differentiated in RDM with 10% FBS to promote maintenance of lamination.

### Retinal Organoid Dissociation and Culturing

ROs were dissociated in 0.5-1mL papain (Worthington Biochemical Corp. #LK003150) in microfuge tubes shaking at 105rpm and at 37°C. Organoids were triturated every hour until larger pieces dissociated (2-4hrs). Dissociated organoids were filtered with a 37μm cell strainer (STEMCELL Technologies #27215) to isolate single cells, incubated in ovomucoid (Worthington), rinsed in Calcium and Magnesium Free Phosphate-Buffered Saline (CMF-PBS), and resuspended in RDM (no FBS). Cells were plated at 50,000 cell/cm^2^. For cultures with a poly-D-lysine substratum, glass-bottom dishes were incubated in 50μg/mL in PBS for 45 min. For Laminin (Sigma #L2020-1MG) substrata, glass-bottom dishes were incubated in 25μg/mL in PBS for 1-2hr. For PDL-LN, PDL was applied first, rinsed, then LN was added. Nitrocellulose (NC) substrata were prepared as Kljavin *et al*. (38). Briefly, a 16cm^2^ piece of NC was dissolved in 15mL methanol, then 3 coats of 100μL was added to each well of a 24-well glass-bottom plate (Cellvis #P24-1.5H-N). Two days later, Neural Cell Adhesion Molecule (EMD Millipore #AG265), N-cadherin (Sino Biological #11039-H03H), Laminin, or PBS control was added to NC-coated wells. For all other dissociated culture experiments, culture dishes were either custom-made 10-15mm diameter glass-bottom dishes [acid-washed coverslip (Fisher Scientific #12-541-B 22×22-1.5) affixed over a hole in a 35mm petri dish (Falcon #351008) or pre-manufactured dishes (Cellvis #D35-10-1.5-N or #D35-20-1.5-N). Culture dishes were flooded with 2mL RDM (no FBS, low phenol red) the day after cells were plated. All hPSC-PR cultures were maintained for 4 days post-dissociation before being fixed or imaged.

### Müller Glia Co-culture

ImM10 immortalized murine MG cells (35) (passage 9-14) were kept in ImM10 Media [Neurobasal Media (Gibco #21103049), 1x Antibiotic-Antimicrobial (Gibco #15240-062), 1x B-27 supplement (Gibco #17504-044), 10% FBS (Peak Serum, Inc. #PS-FBS), 10ng/mL Interferon-gamma (Peprotech #315-05-20UG), and 1x (2mM) Glutamax (Gibco #35050-061)] and passaged with 0.05% Trypsin-EDTA (Gibco #25300054) onto tissue culture petri dishes.

For co-culture with hPSC-derived RO cells, ImM10 cells were passaged as usual and plated at ∼25% confluency on poly-D-lysine coated glass bottom dishes in ImM10 media. The following day, retinal organoids were dissociated as described above and plated on top of the ImM10 dishes, at which point the cells were put in RDM (no FBS, low phenol red). Cultures were kept for 4 days post-dissociation before live confocal imaging and fixation.

### EdU Labeling

On d30, d35, or d38 ROs were treated with 1μM EdU (Click-iT Plus EdU Kit; Life Tech Molecular Probes #C10637) in RDM (2% FBS). At d40, ROs were dissociated and cultured on PDL-LN as described. Dishes were live imaged 4 days post-dissociation, fixed, permeabilized, and blocked (see Immunocytochemistry section for details). Dishes were probed for EdU following manufacturer’s instructions and incubated with 300nM DAPI to stain nuclei. Cells observed for live imaging were subsequently examined for EdU labeling.

### Confocal imaging (live and fixed)

Confocal imaging was performed on a Zeiss LSM 800 inverted microscope. For live imaging, a gas-impermeant chamber was created by affixing a glass ring to the glass-bottom dish (10-20mm diameter) with vacuum grease (Dow Corning #Z273554), filling it with culture media (RDM with no FBS, low phenol red), and sealing with a coverslip. During live imaging, dishes and stage were kept at 37°C with a Zeiss Incubator XLmulti S1. Images were captured with either a Plan Apo 63x oil immersion objective (NA 1.4) or Plan Apo 20x air objective (NA 0.8).

### Multi-Photon Imaging

To stabilize intact, live ROs for multi-photon imaging, on the day before imaging ROs were partially embedded in 3% low-melt agarose (IBI Scientific #IB70056) in PBS by layering 37°C agarose with ROs onto prepared 35mm petri dishes (Falcon #351008) that had previously been filled with Sylgard 184 Silicone Elastomer (Sylgard Material # (204)4019862) and cut to create a conical frustrum to prevent the agarose from detaching. RDM (10% FBS, low phenol red) was added on top to completely submerge the ROs.

Multi-photon imaging was performed at the University of Wisconsin Optical Imaging Core with a Nikon A1R HD upright microscope equipped with dual multi-photon lasers (Coherent Discovery), Piezo insert, and GaAsP detectors. ROs were kept in humidified 37°C chamber with 5% CO_2_ with a Tokei Hit biochamber. 25x APO LWD water immersion objective (NA 1.10) was used for all experiments.

### STED Microscopy

STED microscopy was performed at the University of Wisconsin Optical Imaging Core with a Leica SP8 3X STED confocal microscope with a super Z Piezo stage and HyD detectors. Images were collected with a 100x PLAN APO oil immersion objective (NA 1.4).

### Immunocytochemistry

Cultures were fixed for 15 minutes in 4% paraformaldehyde (Sigma-Aldrich #158127) in Krebs + Sucrose solution, rinsed with PBS, permeabilized for 15 minutes in 0.1% Triton (Sigma #X100-100G) in CMF-PBS, then blocked in 1% Fish Gelatin (Sigma #G7765) + 0.1% Sodium Azide in CMF-PBS. Cultures were incubated in primary antibody (G-actin -1:500 DSHB #JLA20-c, mouse IgM anti-actinA; SV2 – 1:20 DSHB #SV2-s, mouse IgG1 anti-SV2A; Syt1 – 1:200 DSHB #mab48-c, mouse IgG2b) in block for > 1 hour at room temperature or overnight at 4°C. They were then rinsed in block for >30 min at room temperature before incubation in secondary antibody (1:250 Invitrogen #A21-42 AF488 anti-mouse IgM; 1:250 Life Technologies #A11029 AF488 anti-mouse IgG1; 1:250 Invitrogen #A21141 AF488 anti-mouse IgG2b) in blocking solution for > 1 hour at room temperature or overnight at 4°C. Final rinse was done in PBS, with 1:100 phalloidin (Invitrogen #A12379 or #A22287) added to last wash, where used.

### Two-Dimensional Image Analysis

Image analysis was performed using Fiji (ImageJ 1.52p) (57, 58). Where needed for analysis, time-lapse movies were stabilized with the Image Stabilizer Plugin (59). Analysts were masked to condition for all qualitative dynamicity measures (Fig. 2B,E; Fig. 7F). Quantitative dynamicity and displacement rate measurements were made by cropping the time-lapse images around terminals. ImageJ macros scripts were used to threshold on the tdTomato channel to create a mask for each frame. Area of protrusion, retraction, and total terminal, as well as centroid, were measured for each mask to calculate dynamicity and displacement rate (X-Y displacement of the centroid over time). Neurite lengths were calculated by manually tracing a segmented line that was converted to a spline (Fig. 3B, Fig. 4) or a straight line (Fig. 7). Cell Body Displacement Rates and Terminal Displacement Rates (Fig. 3D) were calculated by manually drawing a straight line from the cell body centroid to the terminal centroid at the first frame and last frame of time-lapse series, then capturing the X-Y coordinates of those points. Neurite lengths (Fig. 3E) were calculated as the distance between the cell body centroid and the terminal centroid.

Fluorescence intensity for F-actin and G-actin was measured by thresholding on tdTomato signal, cropping the ROI so that only the terminal was included, then measuring mean intensity. Fluorescence intensity measurements for SV2 and Syt-1 were made by thresholding on tdTomato, cropping the ROI so only the neurite or only the cell body were selected, then measuring mean intensity. Determinations of cell-cell contact were made manually by observing merged time-lapses of the tdTomato and transmitted light channels. Non-PR cells were identified by a lack of tdTomato fluorescence and (for MG) characteristic large, flat, motile morphology. IRM image intensities were measured from maximum projection of time lapse movies. The image intensity of each terminal was subtracted from adjacent background.

### Three-Dimensional Image Analysis

Whole RO time-lapse sequences were manually stabilized in Z dimension by aligning matching slices (using distinct cells as markers) across each timepoint, then running the Image Stabilizer Plugin to stabilize in X-Y dimension. Cells were selected for analysis based on the criteria that their terminal must be distinguishable from nearby cells or debris, the entire cell must be within the imaging section for the duration of the time series, and the cell must be bright enough to assure the terminal and neurite can be distinguished for the duration of the time-lapse. ROIs were made containing cells of interest that were max projected in z to allow two-dimensional visualization of the cell. Two independent analysts (SKR and MJW) reviewed each time-lapse movie, masked and in random order, and marked the cell as having a terminal that is “Dynamic” or “Stationary”. Percent in each category per RO was averaged between analysts. Neurite lengths (Fig. 7G) were calculated by manually tracing a straight line over the axon of a maximum projection of the cell.

### Statistics

Data sets were first tested for normality to determine the appropriate statistical testing. Tests used are listed in the figure legends. All statistical testing was performed with Prism Software (Version 9.1.1 for Windows, GraphPad Software, San Diego, California USA, www.graphpad.com).

## Acknowledgments

We thank Lance Rodenkirch and the University of Wisconsin – Madison Optical Imaging Core (Grant No. 1S10OD025040-01) for use of the upright multiphoton and STED microscopes. This work was supported by grants to T.M.G. from National Institute of Neurological Disorders and Stroke Grant No. 5R01 NS113314-02 and 5R01NS041564 and to D.J.Z. and D.M.G. from National Eye Institute (NEI) Grant No. U01 EY027266-01 and. Funding was also provided by awards to D.M.G by the Retina Research Foundation Emmett Humble Chair, Sarah E. Slack Prevention of Blindness Fund (a component Fund of the Muskingum County Community Foundation), the McPherson Eye Research Institute Sandra Lemke Trout Chair in Eye Research, as well as the Guerrieri Family Foundation to D.J.Z, and Research to Prevent Blindness to D.M.G. and D.J.Z. This study was supported in part by a core grant to the Waisman Center (NICHHD U54 HD090256). S.K.R. was supported by NEI Grant. No. T32 EY027721. A.L.L. was supported by the UW-Madison School of Veterinary Medicine DVM/PhD Program, NEI Grant No. U24 EY029890, and a Kirschstein NRSA Predoctoral Fellowship (NEI Grant No. F30 EY031230).

