## supplemental figures for "Human photoreceptors switch from autonomous axon extension to cell-mediated process pulling during synaptic marker redistribution"

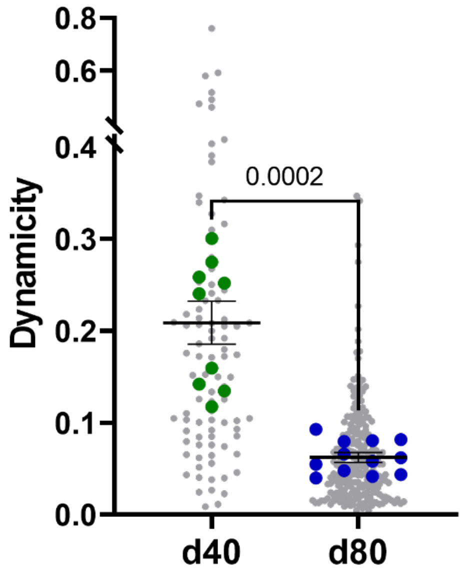


**Supplemental Figure 1. PR terminal dynamics are determined by PR age, not associated RO cell age.** PRs from RO cells dissociated at either d40 or d80 using the WA09 CRX$^{+/tdtomato}$ line were co-cultured with RO cells dissociated at the other age using an unlabeled WA09 line. Dynamicity measurements show a decrease in terminal dynamicity from d40 to d80 regardless of co-cultured RO cell age. P-value shown using Welch's t test.

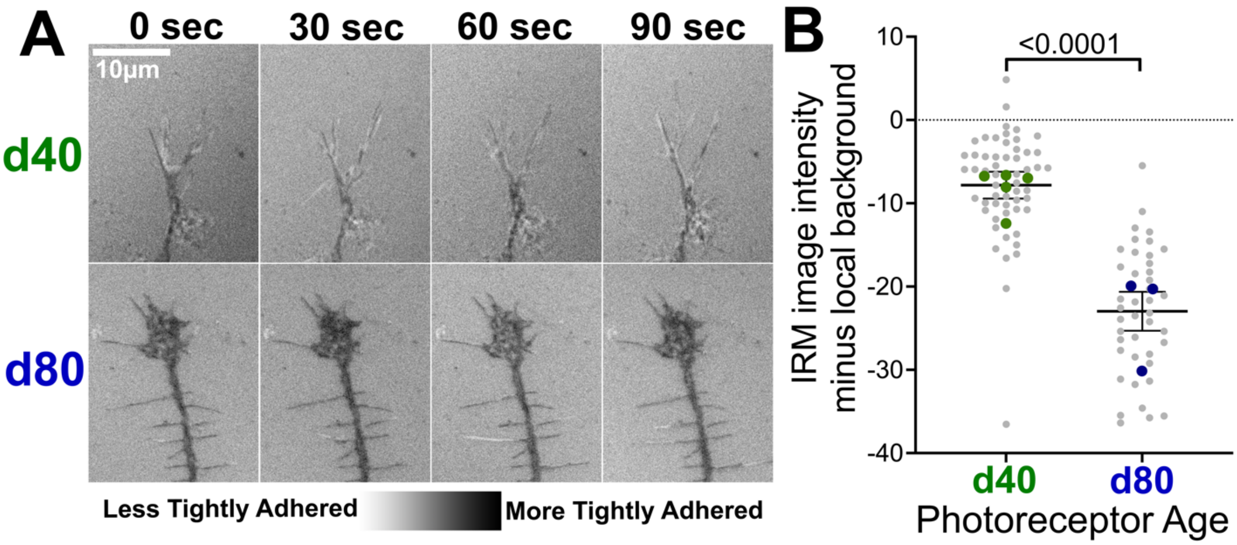


**Supplemental Figure 2. Day 80 PRs are more closely adhered to the substratum than day 40 PRs. A:** Time-lapse interference reflection microscopy (IRM) images of d40 and d80 PR terminals. Dark images suggest that terminals of d80 PRs are more tightly adhered to the substratum than motile d40 PR terminals. D40 terminal IRM images are lighter and more varied over time, which indicates the cell membrane is further from the substratum and more dynamically adhesive. **B:** The intensity of maximum projection images of PR terminals over time was quantified using tdTomato signal to highlight terminals (see methods). The IRM image intensity of d80 terminals was significantly less than d40 terminals P-value shown using Welch's t test.

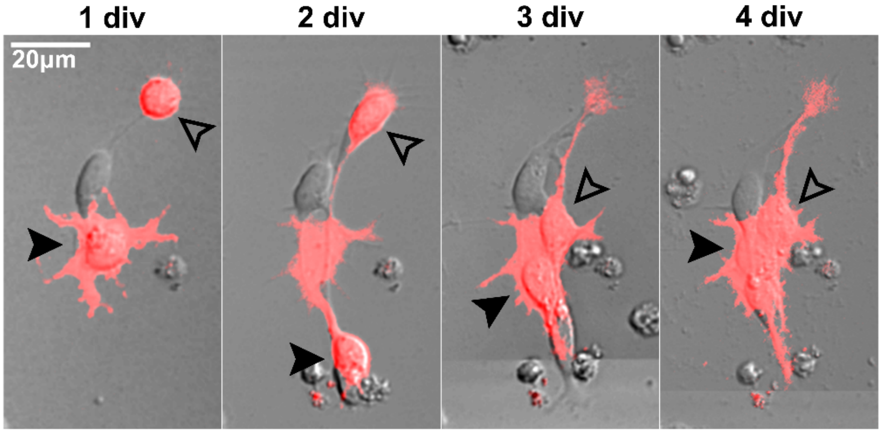


**Supplemental Figure 3. Day 80 PR terminals form when cell body moves away from stationary terminals which can stay in place for days.** Two d80 PRs imaged from 1-4 div both have a terminal formed when the cell body (arrowheads) moves away from its initial position. Note that non-PR cells (tdTomato negative) are in contact with the PRs, including one non-PR that was not present at 1 div.

**Supplemental Movie 1. Day 80 PR co-cultured with MG can undergo dramatic stretching.** 3.25 hours after plating, a d80 PR at the top of the region of interest grows autonomously for a few microns before being dramatically pulled across the region of interest by a motile MG.

**Supplemental Table 1.** Number of biological and technical replicates done for each experiment.

| Figure | Age | Condition | Num. of Biological Replicates (Differentiations) | Num. of Technical Replicates (dishes, wells, or ROs) |
|---|---|---|---|---|
| **1B,C** | d40 | | 3 | 12 |
| | d50 | | 2 | 8 |
| | d60 | | 2 | 6 |
| | d70 | | 2 | 7 |
| | d80 | | 3 | 12 |
| **2B** | d40 (PRs) | | 3 | 12 |
| | d50 (PRs) | | 2 | 8 |
| | d60 (PRs) | | 2 | 6 |
| | d70 (PRs) | | 2 | 7 |
| | d80 (PRs) | | 3 | 12 |
| | d80 (Rods) | | 2 | 7 |
| | d90 (Rods) | | 2 | 4 |
| | d100 (Rods) | | 1 | 3 |
| | d127 (Rods) | | 1 | 4 |
| **2E** | d40 | d30 EdU pulse | 3 | 9 |
| | d40 | d35 EdU pulse | 3 | 9 |
| | d40 | d38 EdU pulse | 3 | 9 |
| **3B** | d40 | | 3 | 12 |
| | d50 | | 2 | 8 |
| | d60 | | 2 | 6 |
| | d70 | | 2 | 7 |
| | d80 | | 3 | 12 |
| **3D,E** | d40-44 | | 4 | 12 |
| | d80-82 | | 3 | 6 |
| **4B** | d40 | glass | 3 | 8 |
| | d40 | PDL | 3 | 8 |
| | d40 | PDL-LN | 3 | 8 |
| | d40 | LN | 3 | 8 |
| | d40 | NC | 3 | 9 |
| | d40 | NC-LN | 3 | 9 |
| | d40 | NC-Ncad | 3 | 7 |
| | d40 | NC-NCAM | 3 | 9 |

| | | | | |
|---|---|---|---|---|
| | d80 | glass | 2 | 6 |
| | d80 | PDL | 2 | 6 |
| | d80 | PDL-LN | 2 | 6 |
| | d80 | LN | 2 | 5 |
| | d80 | NC | 2 | 6 |
| | d80 | NC-LN | 2 | 6 |
| | d80 | NC-Ncad | 2 | 6 |
| | d80 | NC-NCAM | 2 | 6 |
| **5C** | d40 | | 1 | 3 |
| | d50 | | 1 | 4 |
| | d60 | | 1 | 4 |
| | d70 | | 1 | 4 |
| | d80 | | 1 | 4 |
| **5D** | d40 | | 2 | 7 |
| | d50 | | 2 | 8 |
| | d60 | | 2 | 8 |
| | d70 | | 2 | 8 |
| | d80 | | 2 | 8 |
| **6B** | d40-45 | | 2 | 6 |
| | d80-84 | | 2 | 7 |
| **6C** | d40 | | 3 | 12 |
| | d80 | | 3 | 12 |
| **7B** | d40 | | 5 | 18 |
| | d60 | | 3 | 10 |
| | d80 | | 3 | 10 |
| **7C,D,F,G** | d40 | | 5 | 12 |
| | d60 | | 3 | 10 |
| | d80 | | 3 | 10 |
